# Structural analysis of mycobacterial homoserine transacetylases central to methionine biosynthesis reveals druggable active site

**DOI:** 10.1101/808071

**Authors:** Catherine T. Chaton, Emily S. Rodriguez, Robert W. Reed, Jian Li, Cameron W. Kenner, Konstantin V. Korotkov

## Abstract

*Mycobacterium tuberculosis* is the cause of the world’s most deadly infectious disease. Efforts are underway to target the methionine biosynthesis pathway, as it is not part of the host metabolism. The homoserine transacetylase MetX converts L-homoserine to *O*-acetyl-L-homoserine at the committed step of this pathway. In order to facilitate structure-based drug design, we determined the high-resolution crystal structures of three MetX proteins, including *M. tuberculosis* (*Mt*MetX), *Mycolicibacterium abscessus* (*Ma*MetX), and *Mycolicibacterium hassiacum* (*Mh*MetX). A comparison of homoserine transacetylases from other bacterial and fungal species reveals a high degree of structural conservation amongst the enzymes. Utilizing homologous structures with bound cofactors, we analyzed the potential ligandability of MetX. The deep active-site tunnel surrounding the catalytic serine yielded many consensus clusters during mapping, suggesting that *Mt*MetX is highly druggable.

## Introduction

*Mycobacterium tuberculosis* (Mtb) is the causative agent of tuberculosis (TB), a persistent global health threat. In 2017, TB was responsible for the deaths of 1.3 million HIV-negative people and an additional 300,000 deaths among the HIV-positive population.^1,2^ Approximately 10 million people contracted TB that same year.^2^ Despite increased focus since WHO declared TB as a global health emergency back in 1993, progress towards fighting it has been mixed. It ranks among the top ten causes of death among the world’s population and is the single most deadly infectious disease, claiming more lives than HIV/AIDS.^2^ Survivors of TB infections can also suffer significantly reduced quality of life.^3^

The bacilli Calmette-Guerin (BCG) vaccine is capable of protecting children from the most severe forms of TB and is still the preferred method for disease control. However, no vaccine is capable of preventing TB infection in adults either pre or post-exposure.^4^ Existing anti-TB therapy requires treatment of six months or more with a combination of multiple therapeutic agents: rifampicin, ethambutol, isoniazid, and pyrazinamide. Treatment with first-line drugs leads to a cure rate of about 85% of those with drug-susceptible TB strains.^2^ The side effects and length of these treatments has led to compliance issues. This fact, combined with the lack of care in some areas of the world, has spurred the development of multi-drug resistant TB (MDR-TB); 6% of these MDR-TB cases are so antibiotically hardened that they are classified as extensively drug-resistant TB (XDR-TB).^2^ MDR-TB and XDR-TB infections have limited treatment options which are not always successful, particularly in the case of immunocompromised patients and XDR-TB. Current therapeutic regimens are often toxic, require long durations of treatment time, and are estimated to increase treatment costs 8 to 15- fold compared to treating regular TB infections.^5^ This difference is even more extreme in the case of XDR-TB with costs being 25 to 32-times higher. There is a clear need for practical, low-cost TB therapies with orthogonal activity to current antibiotic options.

MetX (previously mis-annotated as MetA in the reference genome of *M. tuberculosis*) is an L-homoserine *O*-acetyltransferase, commonly also referred to as a homoserine transacetylase (HTA) and is a crucial enzyme in the biosynthesis of methionine and threonine (Fig. 1).^6^ It catalyzes the conversion of L-homoserine to *O*-acetyl-L-homoserine (OAHS) by transfer of an acetyl group from Acetyl-CoA to the γ-hydroxyl of homoserine.^7^ OAHS is an essential precursor to methionine as well as other bacterial metabolites, such as *S*-Adenosyl-L-methionine (SAM). The modification of L-homoserine is a committed step in methionine and SAM synthesis.

**Figure 1.**
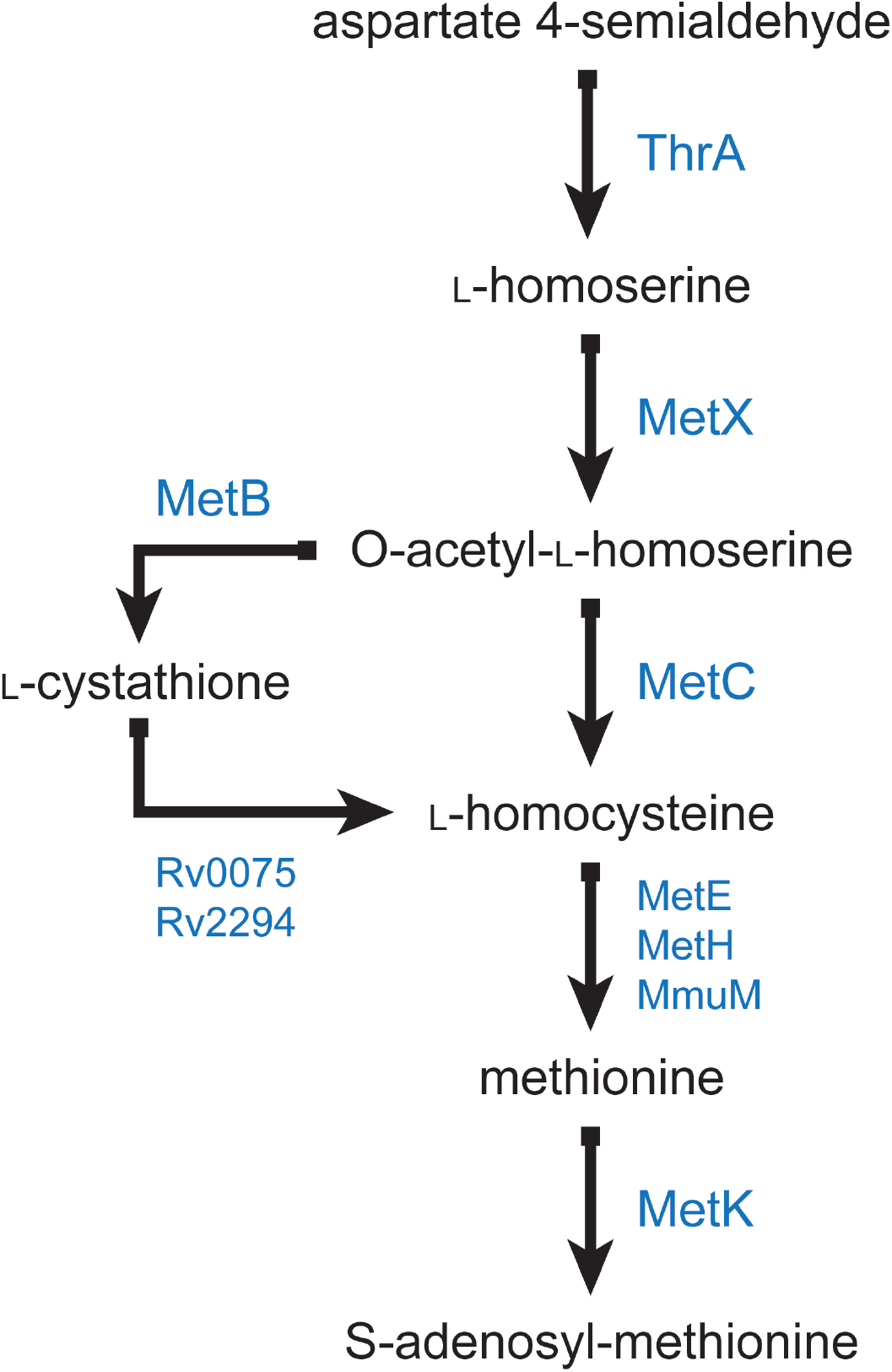
Overview of the *M. tuberculosis* biosynthesis pathways dependent on MetX converting L-homoserine into either *O*-succinyl-L-homoserine or *O*-acetyl-homoserine. Inhibition of MetX will affect not only methionine synthesis but also the production of SAM and threonine, among other necessary metabolites.

Bacterial studies have shown that deletion of *met2* gene, which encodes an analogous HTA, is lethal without methionine supplementation.^8,9^ Further studies using immunocompetent and immunocompromised mice demonstrate that deletion of *metX* generates auxotrophic mutants unable to establish infection.^10^ If starved of threonine and methionine *in vitro*, Mtb Δ*metX* dies quickly. Furthermore, Δ*metX* mutant strain is unable to proliferate inside of human macrophages. Also, it has recently been demonstrated that *metX* is required for maintaining bacterial survival during chronic Mtb infection.^11^ Together, these data suggest that Mtb is unable to scavenge biosynthetic intermediates from the host for methionine synthesis, making for a uniquely exploitable vulnerability for the development of antibacterial agents.^12^

Efforts are already underway to inhibit HTA in *Cryptococcus neoformans* for use as an antifungal agent.^8^ Targeting of a similar pathway for aspartate production has already yielded some promising selective inhibitors for *Streptococcus pneumoniae* and *Vibrio cholerae*.^13^ A similar structurally guided approach to discovering specific inhibitors is an attractive alternative to traditional antibiotic killing of MDR-TB and XDR-TB.

Here we report the structures of three homologous MetX enzymes from *Mycobacterium tuberculosis* (*Mt*MetX), *Mycolicibacterium abscessus* (*Ma*MetX), and *Mycolicibacterium hassiacum* (*Mh*MetX) and compare them to previously solved structures of HTAs in order to begin development of selective inhibitors of *Mt*MetX via a structurally based approach. Using the *Mt*MetX structure as a guide, we also elucidate the druggability of the enzyme and propose that it is an excellent candidate for small molecule drug discovery.

## Results

### Overview of mycobacterial MetX structures

The three solved MetX structures include residues 15–70, 77–372 from *Mh*MetX (Fig. 2A), 10– 379 from *MaMetX* (Fig. 2B), and 7–372 from *Mt*MetX (Fig. 2C). Two copies of each monomer exist in the asymmetric unit of all three structures. MetX can be divided into two distinct structural domains, the catalytic domain, and the lid domain.

**Figure 2.**
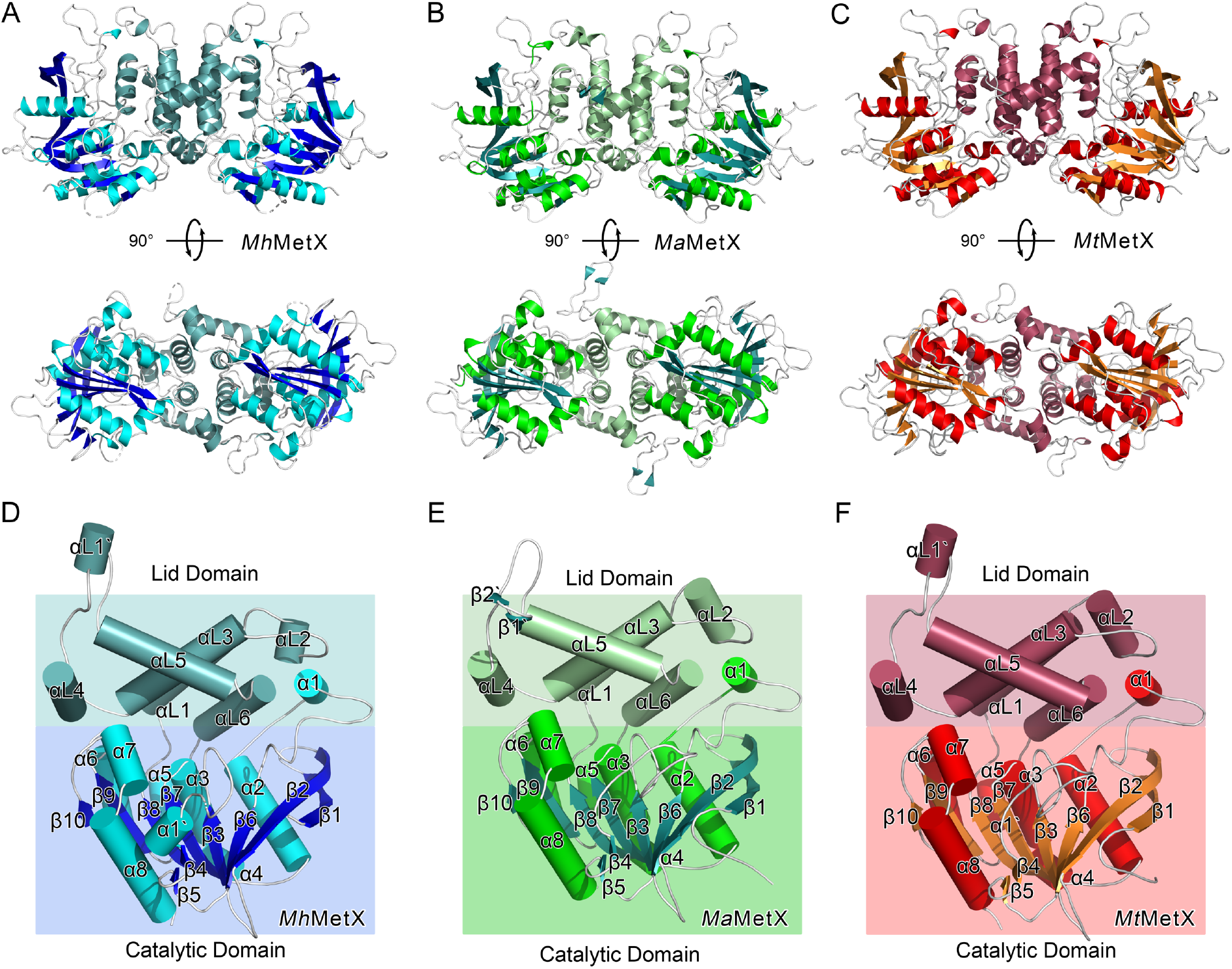
(A) Ribbon diagram of *Mh*MetX. Catalytic domain sheets are shown in blue, helices of the catalytic domain are in cyan, and the lid domain secondary structure is shown in blue-green. (B) Ribbon diagram of *Ma*MetX. Catalytic domain sheets are shown in green-blue, helices in bright green, and the lid domain in mint. (C) Ribbon diagram of *Mt*MetX. Catalytic domain sheets are shown in orange, helices in red, and the lid domain in burgundy. (D-F) Simplified cartoon structures of *Mh*MetX, *Ma*MetX, and *Mt*MetX monomers with topological labels and both domains.

The organization of the catalytic domains’ fold marks MetX as members of the α/β-hydrolase super-family. It is a highly diverse family that includes proteases, lipases, and esterases, among many others.^14–16^ A canonical 8-stranded β-sheet fold with twisted, parallel topology forms the core of α/β-hydrolases.^17^ Several α-helices flank either face of this fold, though their number and location are different depending on the specific protein. The catalytic domain comprises residues 15–181, 297–372 of *Mh*MetX (Fig. 2D), residues 17–183, 304–379 of *Ma*MetX (Fig. 2E) and residues 17–181, 311–372 of MtMetX (Fig. 2F). The catalytic domain contains the active site tunnel with its a canonical catalytic triad.

Assembly occurs at an anti-parallel four-helix bundle motif (αL1 and αL3) in the lid domain. The total interface area of the dimer is ~1700 Å^2^.^18^ Two additional helices help to strengthen the interaction with hydrogen bonds and van der Waals contacts (αL4 and αL5). Like other previously studied HTAs, MetX forms solution dimers at this interface. These dimers have been shown to be physiologically relevant in other HTAs and are likely important for MetX as well.^19^ The lid domain comprises residues 184–285 of both *Mh*MetX and *Mt*MetX, and residues 187– 292 of *Ma*MetX, between β8 and α5 (Fig. 3A).

**Figure 3.**
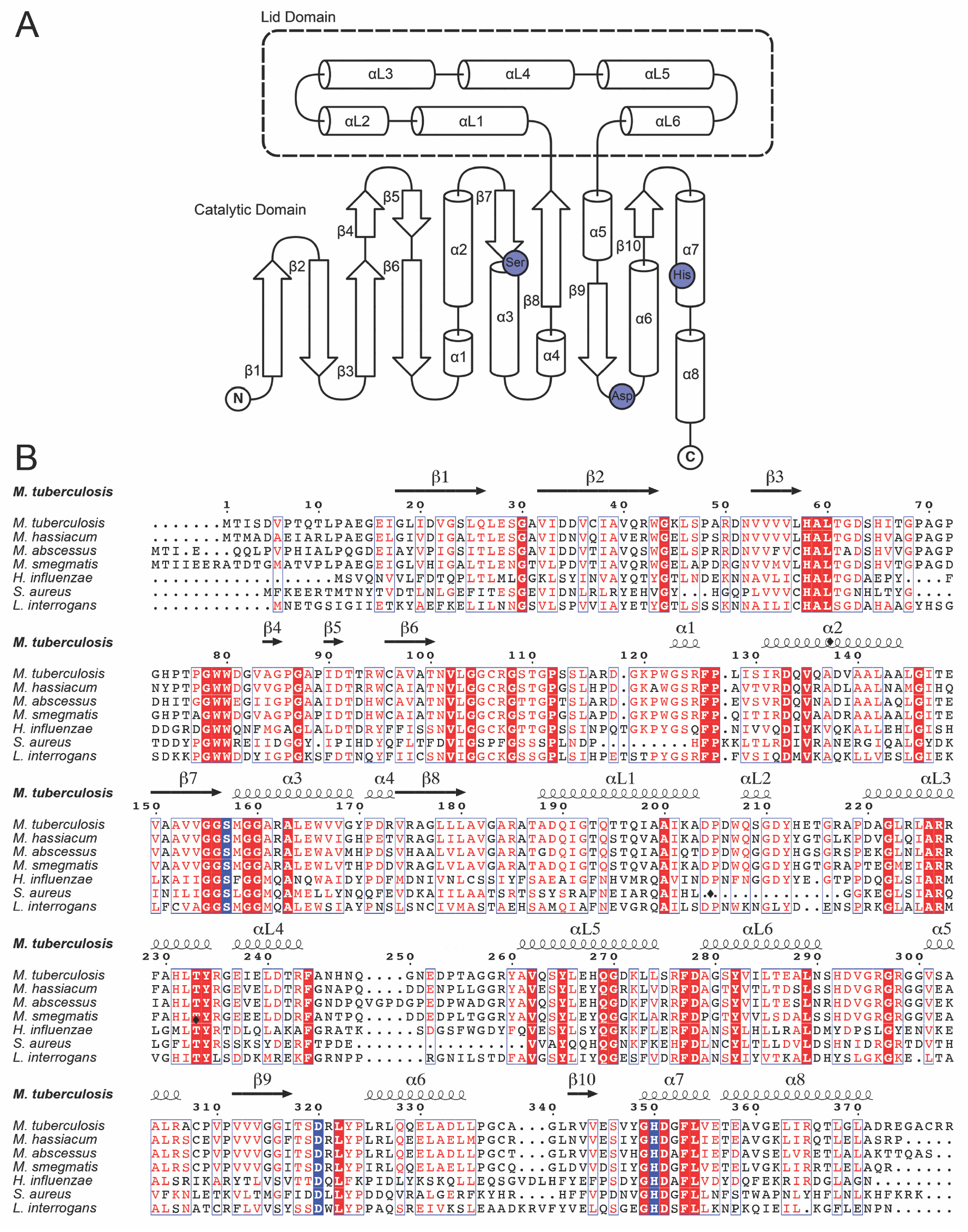
(A) Simplified schematic diagram of conserved secondary structural elements of MetX. (B) Sequence alignments of *Mh*MetX, *Ma*MetX, *Mt*MetX, *Hi*HTA, *Sa*HTA, and *Li*HTA with assigned secondary structure denoted above. Catalytic residues are marked with diamonds.

The space between the catalytic and lid domains forms a deep active-site tunnel. At the end of this tunnel sits the nucleophilic serine residue. The tunnel is lined with polar residues highly conserved among other known HTA structures. Thr61/61/64, Arg227/227/230, Tyr234/234/237, and Asp351/351/358 for *Mh*MetX, *Mt*MetX, and *Ma*MetX respectively all surround the active site to help facilitate the binding of acetyl-CoA and homoserine (Fig 3B).^20^

### Catalytic Mechanism

The catalytic triad of Nucleophile-His-acid is the α/β-hydrolase family’s most conserved feature. Just as in other known HTA structures, *Mt*HTA, *Mh*HTA, and *Ma*HTA contain a serine, aspartic acid, and histidine in the active site. HTAs have a serine between β7 and α3, an aspartic acid on the loop between β9 and α6, and histidine on α7 for these residues. For *Mt*HTA and *Mh*HTA, Ser157, Asp320, and His350 comprise the active site; *Ma*HTA’s triad is comprised of Ser160, Asp327, His357 (Fig. 4A). The catalytic serine sits at the end of a deep catalytic tunnel (Fig 4B).

**Figure 4.**
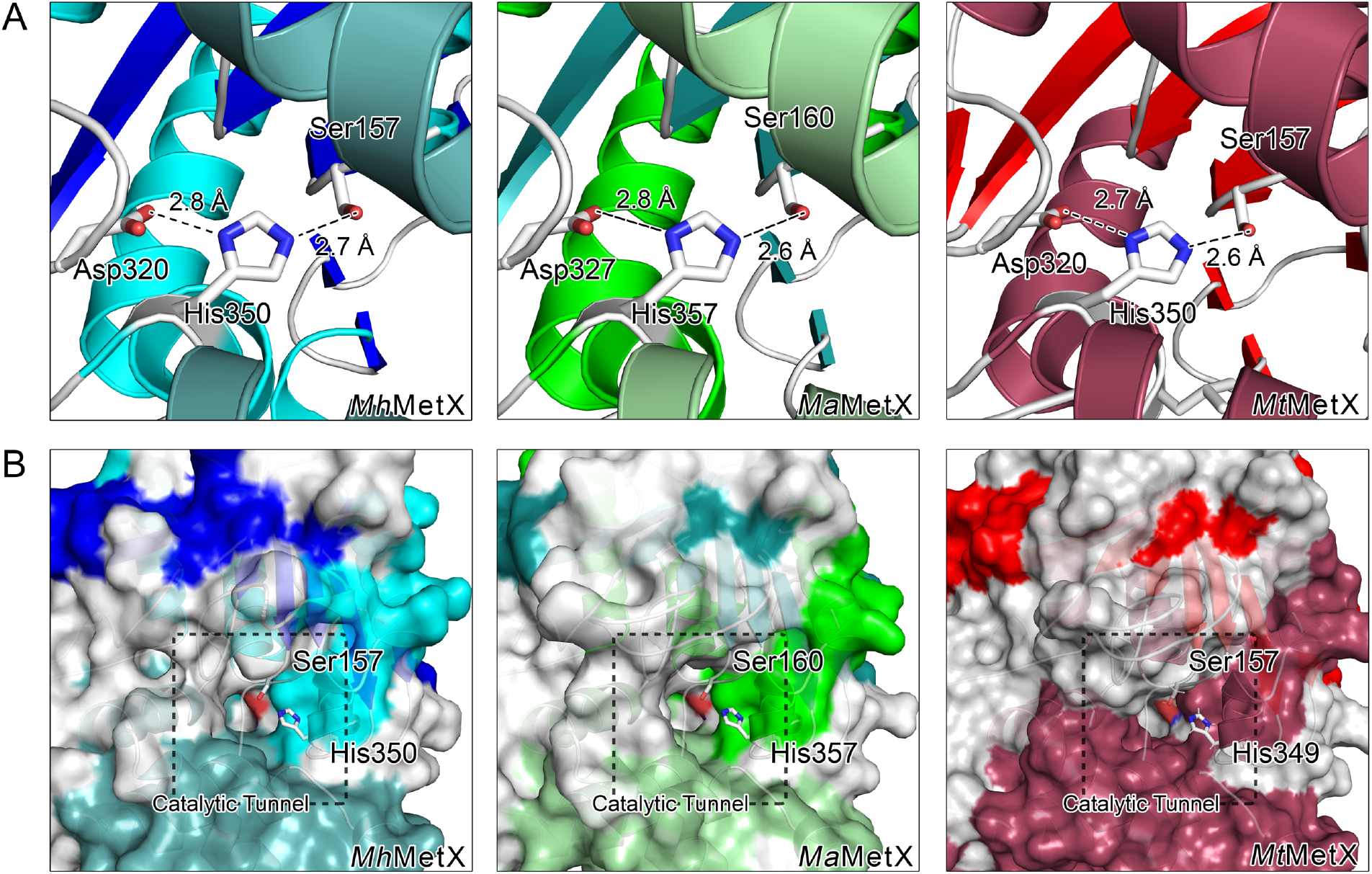
(A) Ribbon diagrams of *Mh*MetX, *Ma*MetX, and MtMetX active site residues. Catalytic residues are shown in stick representation with key hydrogen bonding distances between heavy atoms. (B) Cross-sectional diagrams of *Mh*MetX, *Ma*MetX, and MtMetX catalytic tunnels with the catalytic serine and histidine residues highlighted.

Studies in *H. influenzae* and *Schizosaccharomyces pombe* suggest that the mechanism of HTAs is based on a “ping-pong” reaction.^21^ In the proposed mechanism, acetate is first transferred to serine from acetyl-CoA in the “ping” step to create an acetyl-enzyme intermediate through an attack of the acetyl-CoA thioester bond. In the “pong,” L-homoserine breaks down a tetrahedral intermediate of the acetyl-enzyme to complete the transfer to OAH.^20^ Histidine and aspartic acid function as bases to activate the serine residue for attack and can also most likely assist in deprotonation of the L-homoserine.^21^

The active site residues of *Mt*MetX, *Mh*MetX, *Ma*MetX, and *Ms*HTA have similar geometry to other bacterial and fungal HTAs. Just as in other structures, the serine sits in a strained conformation at the end of the active site tunnel, ideally positioned for its function as the nucleophile. The histidine and aspartic acid residues are within hydrogen-bonding distance for activation of the serine (Fig. 4). The active sites of *Li*HTA and *Sa*HTA show the most deviation. In the structure, His344 of *Li*HTA exists in two different conformations with equivalent occupancy.^22^ In the *Sa*HTA structure, His296 is more disordered with a high B-factor (B = 72 Å^2^) for its imidazole ring.^23^ Temperature factors of the equivalent rings in *Mh*MetX is B = 19 Å^2^, while both *Mt*MetX and *Ma*MetX are B = 21 Å^2^, values very similar to the *Hi*HTA structure with B = 16 Å^2^.

### Structural Comparisons to Other HTAs

The overall fold is similar to previously solved HTAs, specifically *Ms*HTA (PDB entry 6IOG)^24^, *Hi*HTA (PDB entry 2B61)^25^, *Sa*HTA (PDB entry 4QLO)^23^, *Li*HTA (PDB entry 2PL5)^22^. The three solved MetXs are most similar in length to *Ms*HTA (374 residues), with *Li*HTA (366 residues) and *Sa*HTA (322 residues) being smaller proteins than MetX and *Hi*HTA (377 residues) being slightly longer. The significant differences lie in the loop lengths and a few short secondary structural elements. The most notable secondary structure difference occurs in the length of β1 and β2, which are both extended in the *Mt*MetX, *Mh*MetX, *Ma*MetX, and *Ms*HTA structures when compared to the other related HTA structures. In *Li*HTA, *Hi*HTA, and *Sa*HTA, these sheets are both subdivided by loops, whereas they are continuous in *Mt*MetX, *Mh*MetX, *Ma*MetX, and *Ms*HTA. The extended loop between β3 and β4 show differences in secondary structure content, length, and orientation. Both *Mh*MetX and *Hi*HTA contain a short sequence with helical propensity adjacent to β4. *Ma*HTA and *Li*HTA have no secondary structure as assigned by DSSP in this same region^26,27^, while *Sa*HTA is unique in substituting β4 and β5 for a longer helix.

As the lid domains are the least conserved among the six HTA structures and within the α/β- hydrolase family as a whole, it is not surprising that this region has some of the most substantial structural deviations. The loop between αL4 and αL5 appears unique in each structure. *Mt*MetX and *Ms*HTA both contain a small helical region (αL1’); *Ma*HTA features a unique pair of short anti-parallel sheets (β1’, β2’). *Hi*HTA merely contains an unstructured loop, while *Sa*HTA omits the majority of the residues entirely, with only a short linker between αL4 and αL5. *Li*HTA is perhaps the most variant, as its loop affects the length and orientation of αL5. Due to the high degree of variability, this loop is likely not critical for MetX’s function or assembly.

The polar residues which line the active site tunnel show conservation between variants of HTA. Additionally, the motifs surrounding each are nearly identical. Notably, Thr61 (substituted for Ser in *Li*HTA) is in the middle of a HAL**T**GD motif, and Asp351 sits next to the conserved region, adjacent to the catalytic His, in a GH**D**(G/A)FL motif. While the lid domain contains a much lower amount of structural conservation globally, the stretch of highest convergence appears in αL3, which contains both Arg227 and Tyr234 and directly forms the other side of the catalytic tunnel. αL1, the other partner in the four-helix bundle with αL3, also shows a fair degree of sequence conservation. Interestingly, *Sa*HTA stands as an outlier when comparing tunnel dimensions, being much narrower and more restricted when compared to the other four HTA structures.

Alignment of *Mt*MetX, *Ma*MetX, and *Mh*HTA structures using the FATCAT algorithm^3^ demonstrate their high degree of structural similarity (Table 2). Across the examined monomers, all are significantly similar to one another (*P* < 0.05). Structural alignment RMSDs ranged from 0.52–3.04, and sequence similarity was in the range of 38.2–88.5%. *Mh*MetX, *Ma*MetX, and *Mt*MetX ranked in the list of closest structural neighbors currently available on the PDB when applying FATCAT to all related structures. Interestingly, *Ms*HTA is as good if not better an approximation to *Mt*MetX as either *Mh*MetX or *Ma*MetX, suggesting that it may function as a good analog in assays where *Mt*MetX cannot be directly utilized. However, when assaying *Mt*MetX to determine its viability for crystallography, the T_m_ was found to be between 40–41 °C by differential scanning fluorimetry (DSF) between pH 6.9–8.5 in Tris-HCl. Because of *Mt*MetX’s relative thermal stability under physiological conditions, we have chosen to focus on *Mt*MetX when investigating future drug development.

### Potential Ligandability of MetX

Druggability of a protein is understood to be a measure of the relative ease of developing a small molecule, which will effectively modulate its activity *in vivo.*^28,29^ Druggability depends on the pharmacodynamics and pharmacokinetics of the host and the pathogen making it difficult to predict computationally. Ligandability is a necessary condition for druggability, but is a more easily quantified metric for the development of inhibitors.^30^ Understanding the ligandability is a critical first step before embarking on drug discovery. An estimated 60% of small-molecule searches fail due to the target site not being sufficiently druggable, which is positively correlated with not being sufficiently ligandable.^31,32^ The availability of high-resolution structures for MetX opens up the possibility of performing direct therapeutic target discovery, provided it proves to be sufficiently ligandable.

The fundamental principle of drug discovery is that biologically active ligands are complimentary in molecular features and shape to the receptor. These can include physiochemical properties such as hydrophobicity, size, as well as the enclosure of the binding pocket and its promiscuity.^33,34^ A strictly predominantly hydrophobic pocket might indicate a promiscuous binding site that may accommodate a wide-varying of ligands in different modes, some hydrophobic patches are still nevertheless ideal for ligand design. All three solved MetX have a similar, conserved hydrophobic patch (Fig. 5A-C) running along the inside of the active site tunnel. Binding models for homoserine (Fig. 5D) and acetyl-CoA (Fig. 5E) were created by hybridizing the *Mt*MetX structures with available *Ms*HTA structures that have been co-crystallized with both substrates^24^ in order to better understand their coordination within the binding pocket. While the hydrophobic patch does not appear critical to the recognition of either, its proximity to the catalytic site residues make it an ideal feature to leverage for designing hydrophobic ringed small molecules with flexible tail groups that could orient inside of the cleft similar to acetyl-CoA.

**Figure 5.**
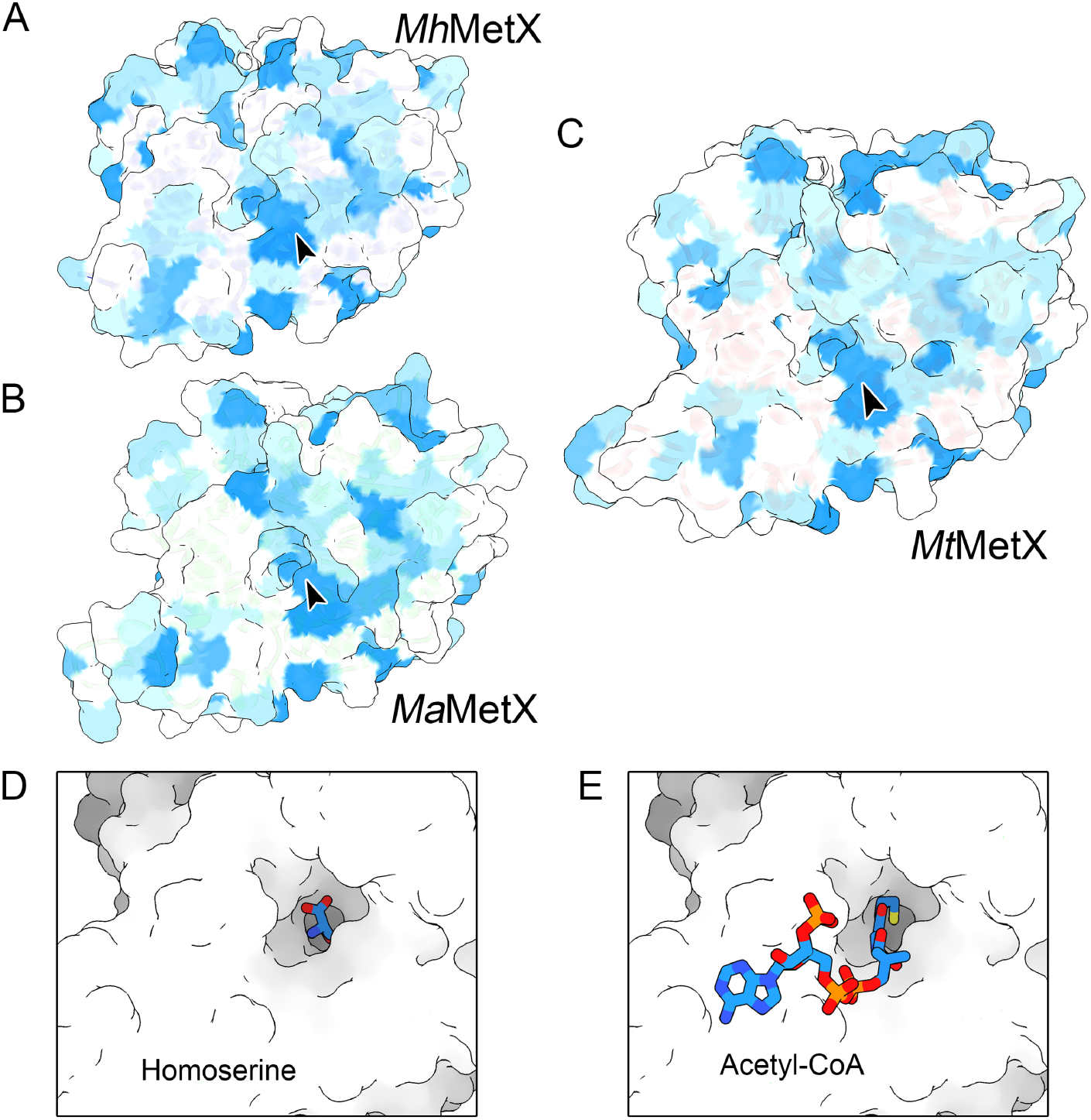
(A-C) Hydrophobicity of surface residues on the Kyte and Doolittle scale of *Mh*MetX, *Ma*MetX, and *Mt*MetX (blue hydrophobic; white hydrophilic). A hydrophobic patch extends from the apex of the active site pocket to the surface of the protein (arrow), which may be an exploitable feature when designing small molecule inhibitors. (D) L-homoserine binding hybrid binding model created using a previously solved *Ms*HTA•HSE structure (PDB ID 6IOH) by aligning the monomer backbone with *Mt*MetX followed by energy minimization. (E) Acetyl-CoA binding model created from the apo *Mt*MetX and *Ms*HTA•acetyl-CoA (PDB ID 6IOI) structures with energy minimization.

In order to further understand the active site’s ligandability, FTMap was utilized.^35–37^ The FTMap algorithm uses a set of small organic probes to sample a protein surface for binding hotspots computationally. Each organic probe is first rigidly docked before being energy minimized.

Areas on the protein’s surface where multiple probes bind are clusters. Binning these clusters based on their member’s average free energy yields a consensus site (CS), a location on the protein’s surface where small molecules are likely to bind. A CS strength (S) is defined as the number of probe clusters within the consensus cluster. A cluster of S > 16 represents a site targetable by a ligand.^36,37^ A second CS should be located somewhere within 8 Å of the primary cluster. Of the eleven CS identified by FTMap, eight reside somewhere within the active site tunnel (Fig. 6). The closest CS near the catalytic Ser have S = 19, S = 16 and S = 13. One CS with S = 13 forms across from the active site on the lid domain, but no other high strength CS exists near it, making it an unlikely site for drug binding. *Ma*MetX shows a similar pattern with the top clusters appearing nearly overtop those of *Mt*MetX with S = 26, S = 13 and S = 10. These results, when overlaid with the hybrid substrate-binding models, suggest that MetX is ligandable.

**Figure 6.**
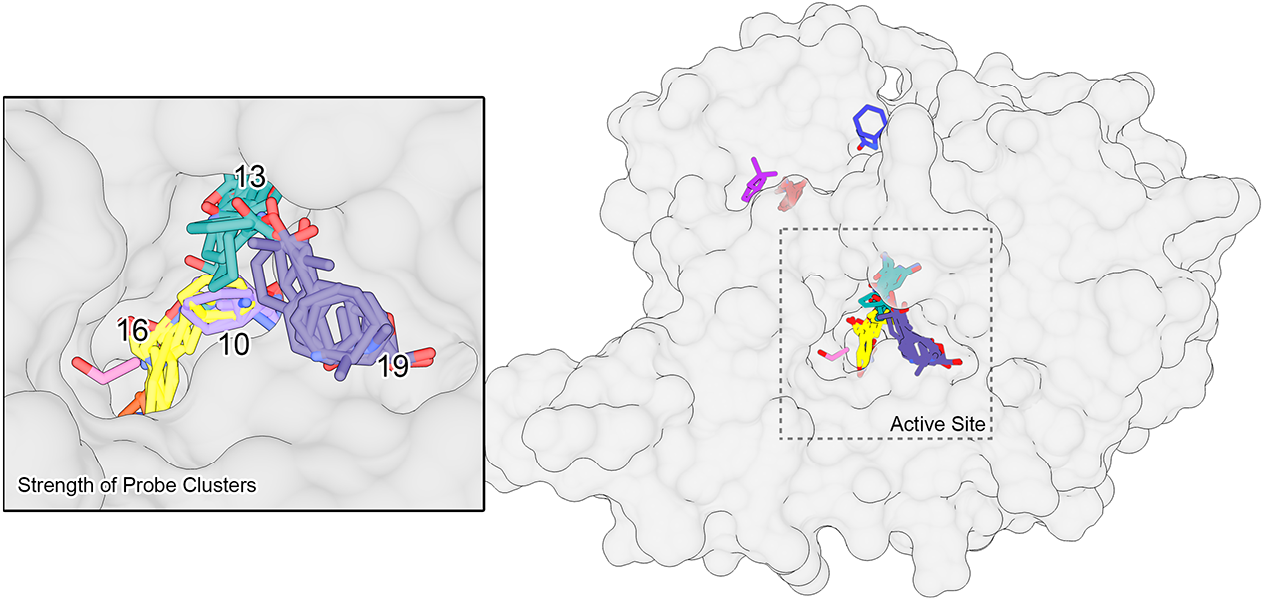
Probe clusters generated from the analysis of monomeric *Mt*MetX using FTMap to evaluate druggability. A few of the critical cluster strengths, the number of probes found in each cluster, are shown in the active site inset.

## Discussion

All three MetX structures show a high degree of overall similarity to previously studied HTAs from both bacteria and fungi. Differences in enzyme size are accounted for by the length of loop regions. Differences in secondary structural elements arise primarily from variations within these loops. All three solved structures crystallized as dimers at the expected interface and orientation that corresponds to predicted physiologically active assembly.

In addition to sharing the overall domain and fold arrangement, the conserved location of the Ser-His-Asp catalytic triad’s position within the active site tunnel may help protect the enzyme from covalent modification and deactivation. Engineered β-lactones with hydrophobic tails have already been shown to inhibit the activity of *Hi*HTA *in vitro* through the formation of adducts. There may be therapeutic value in modifying their structure to enhance specificity towards Mtb.^38^ However, their lack of *in vivo* inhibition of *Hi*HTA suggests that a more efficient method for disrupting methionine biosynthesis lies in small molecule inhibitors. These would be less prone to bacterial inactivation and less prone to the exclusion by Mtb’s complex lipid cell wall. The high degree of structural similarity with previously solved homologs provides an excellent foundation for *in silico* compound screening and structure driven drug design methodologies.

The FTMap cluster data also provides aid towards a Fragment-Based Drug Discovery (FBDD). Previous research has shown that promising core fragments typically bind in the highest strength CS.^39^ While the fragment molecules are too small on their own to have a useful affinity, neighboring CS probe structures can be then be linked to the core fragment to build up a high-affinity ligand. Furthermore, virtually-linked fragments could be screened *in silico* against existing chemical homologs using a tool such as ROCs.^40^

*M. abscessus* MetX was initially chosen for study due to its high similarity to the Mtb variant. However, the structure and CS data argue that it might make an excellent secondary target for drug development. Many compounds that inhibit *Mt*MetX are also likely to affect *Ma*MetX. *M. abscessus* is an emerging public-health threat, primarily implicated in pulmonary infections.^41^ Cross-species gene transfer has helped to create multidrug-resistant strains, some even showing resistance to TB drugs such as rifampin.^42,43^

In summary, by reporting the first medically relevant *Mt*MetX and *Ma*MetX crystal structures, we hope that new avenues of structure-based drug design will be open for developing targeted and effective therapeutics.

## Experimental procedures

### Protein Expression and Purification

Constructs of *Mt*MetX, *Mh*MetX, and *Ma*MetX were prepared from polymerase chain reaction (PCR) products amplified from corresponding genomic DNA and subcloned into a pCDF-NT vector. Following primers were used for amplification:

metXmha_Nco GACACCATGGCCGAAGGCGAACTCG,
metXmha_Hind CTGAAGCTTATGACGCCAACTCCAACGTC,
metXmab_Nco GAGACCATGGCTCTACCCCAGGGCGATGAG,
metXmab_Hind CTCAAGCTTACTTGGCCAGTGCGAGCG,
metXmtb_BspH GAGATCATGACGCTGCCCGCCGAAG,
metXmtb_Hind CACAAGCTTAATCAGCCAATCCCAGTGTCTG.

pCDF-NT is a modified pCDF-Duet1 plasmid (Novagen) encoding His_6_ tag followed by a tobacco etch virus (TEV) protease cleavage site. The pCDG-NT:His_6_-MetX plasmids were transformed into *Escherichia coli* Rosetta (DE3) competent cells (Novagen). Constructs were grown in LB to an OD_600_ of 0.6 at 37°C in the presence of streptomycin and chloramphenicol. Cultures were then cooled in an ice bath to 18°C before the addition of 200 µM Isopropyl β-D-thiogalactopyranoside (IPTG) and 2% v/v ethanol. After inducing overnight for 16 hours, cultures were centrifuged at 5,000 rpm and lysed through two passes through a Microfluidizer (Microfluidics). Cell debris was removed via centrifugation at 18,000 rpm for 1 hour at 4°C. Protein was purified by passage over a Ni-affinity column containing His-Trap chelating resin from GE Healthcare Life Sciences. The column was washed with a buffer containing 20 mM Tris pH 8.0, 300 mM NaCl, and 10 mM imidazole. The protein was then eluted using the same buffer with 250 mM imidazole. The elution fraction was dialyzed overnight at 4°C into 20 mM Tris pH 8.0, 150 mM NaCl alongside TEV protease to release the His_6_ tag. The passage of the protein back over the same His-Trap column removed the TEV protease, and tag before a polishing pass was performed over a Superdex 200 gel filtration column (GE Healthcare Life Sciences) in 20 mM Tris 7.5, 100 mM NaCl buffer.

### *Crystallization of* Mt*MetX*, Mh*MetX, and* Ma*MetX*

Initial crystallization screens were performed using the MCSG (Anatrace) and JCSG (Qiagen) crystallization suites on a Mosquito (TTP Labtech) in vapor-diffusion hanging-drop 96-well format plates. Each drop was created from 1 µL of protein solution and 1 µL well solution. Plates were allowed to grow at 18°C and monitored daily. Initial *Mh*MetX crystals were obtained in 0.2 M Ca acetate, 20% PEG3350. These crystals had spherulite-like morphology. Using Additive Screen (Hampton Research), the optimized prism-like crystals were obtained in 0.15 M Ca acetate, 21% PEG3350, 3% 1,6-diaminohexane, 0.05 M CHES pH 9.5. Initial *Ma*MetX crystals were obtained in 1.2 M NaH_2_PO_4_, 0.8 M K_2_HPO_4_, 0.2 M Li sulphate, 0.1 M CAPS pH 10.5. A pH grid optimization led to final crystallization solution: 1.2 M NaH_2_PO_4_, 0.8 M K_2_HPO_4_, 0.2 M Li sulphate, 0.1 M CHES pH 9.0.The *Mt*MetX crystal was harvested directly from the MCSG screen with crystallization solution 0.1 M Tris-HCl, pH 8.5, 1.8 M magnesium sulfate. Each tray was immediately set up after concentrating fractions from the Superdex 200 to 13.8 mg/mL. The crystal from which the data set was derived grew within 48 hours.

### Data and Structural Determination

Crystals of each MetX construct were flash-frozen in liquid nitrogen after cryoprotection by transfer into the corresponding crystallization solutions supplemented with 20-25% glycerol. Data from *Mh*MetX and *Ma*MetX crystals were collected at the Stanford Synchrotron Radiation Lightsource beamline 9-2 using a Dectris Pilatus 6M detector at 1.000 Å wavelength. Data from *Mt*MetX crystal were collected at the SER-CAT beamline (22-ID) at the Advanced Photon Source using an Eiger 16M detector at 1.00 0 Å wavelength. Data were indexed, integrated, and scaled using *XDS* and *XSCALE*.^44^

The structure of *Mh*MetX was determined by molecular replacement using Phaser ^45^ and the structure of homoserine *O*-acetyltransferase from *Bacillus anthracis* (PDB ID 3I1I) as a search model. The structure of *Ma*MetX was determined by molecular replacement using the *Mh*MetX structure as a search model. The crystal structure of *Mt*MetX was determined using molecular replacement in Phaser ^45^ using the *Ma*MetX structure as a search model. Model building was performed using Coot^46^; iterative refinement was done via *phenix.refine*.^47,48^ Data and refinement statistics are summarized in Table 1.

**Table 1.** Data collection and refinement statistics.

|  | *MhMetX*<sup>a</sup><br>PDB ID 5W8O | *MaMetX*<br>PDB ID 5W8P | *MtMetX*<br>PDB ID 6PUX |
| --- | --- | --- | --- |
| **Data collection** |  |  |  |
| Space group | *P*2<sub>1</sub> | *P*6<sub>5</sub> | *P*2<sub>1</sub>2<sub>1</sub>2<sub>1</sub> |
| Cell dimensions: |  |  |  |
| *a*, *b*, *c* (Å) | 47.75, 93.23, 81.59 | 197.92, 197.92,<br>51.30 | 47.46, 86.68, 208.77 |
| $\alpha$, $\beta$, $\gamma$ (°) | 90, 94.78, 90 | 90, 90, 120 | 90, 90, 90 |
| Resolution (Å) | 47.58–1.47 (1.51–<br>1.47)<sup>b</sup> | 49.48–1.69 (1.73–<br>1.69) | 44.71–1.90 (1.95–<br>1.90) |
| *R*<sub>sym</sub> | 0.040 (1.013) | 0.124 (0.963) | 0.143 (1.361) |
| *CC*<sub>1/2</sub><sup>c</sup> | 1.0 (0.695) | 0.994 (0.678) | 0.997 (0.672) |
| *I* / $\sigma$(*I*) | 18.01 (1.46) | 8.70 (1.78) | 11.21 (1.29) |
| Completeness (%) | 98.3 (99.1) | 99.3 (98.9) | 99.8 (99.7) |
| Redundancy | 3.8 (3.8) | 5.6 (5.8) | 7.4 (7.3) |
| **Refinement** |  |  |  |
| Resolution (Å) | 47.58–1.47 | 49.48–1.69 | 44.71–1.90 |
| No. reflections (total<br>/ free) | 118,621 / 5,840 | 128,098 / 6,439 | 68,843 / 3,463 |
| *R*<sub>work</sub> / *R*<sub>free</sub> | 0.153 / 0.183 | 0.158 / 0.174 | 0.168 / 0.202 |
| No. atoms: |  |  |  |
| Protein | 5,156 | 5,488 | 5,390 |
| Ligand/ion | 12 | 44 | 60 |
| Water | 593 | 787 | 514 |
| *B*-factors: |  |  |  |
| Protein | 25.2 | 20.9 | 32.4 |
| Ligand/ion | 36.3 | 48.2 | 69.0 |
| Water | 36.7 | 34.8 | 39.0 |
| All atoms | 26.4 | 22.8 | 33.3 |
| Wilson *B* | 19.6 | 18.9 | 31.3 |
| R.m.s. deviations: |  |  |  |
| Bond lengths (Å) | 0.005 | 0.006 | 0.003 |
| Bond angles (°) | 0.765 | 0.789 | 0.572 |
| Ramachandran<br>distribution (%): |  |  |  |
| Favored | 98.37 | 97.96 | 97.66 |
| Allowed | 1.63 | 2.04 | 2.34 |
| Outliers | 0 | 0 | 0 |

<sup>a</sup>All data were collected from single crystals.<sup>b</sup>Values in parentheses are for the highest-resolution shell.<sup>c</sup>*CC*<sub>1/2</sub> correlation coefficient between intensities from two random half-data sets.

**Table 2.**
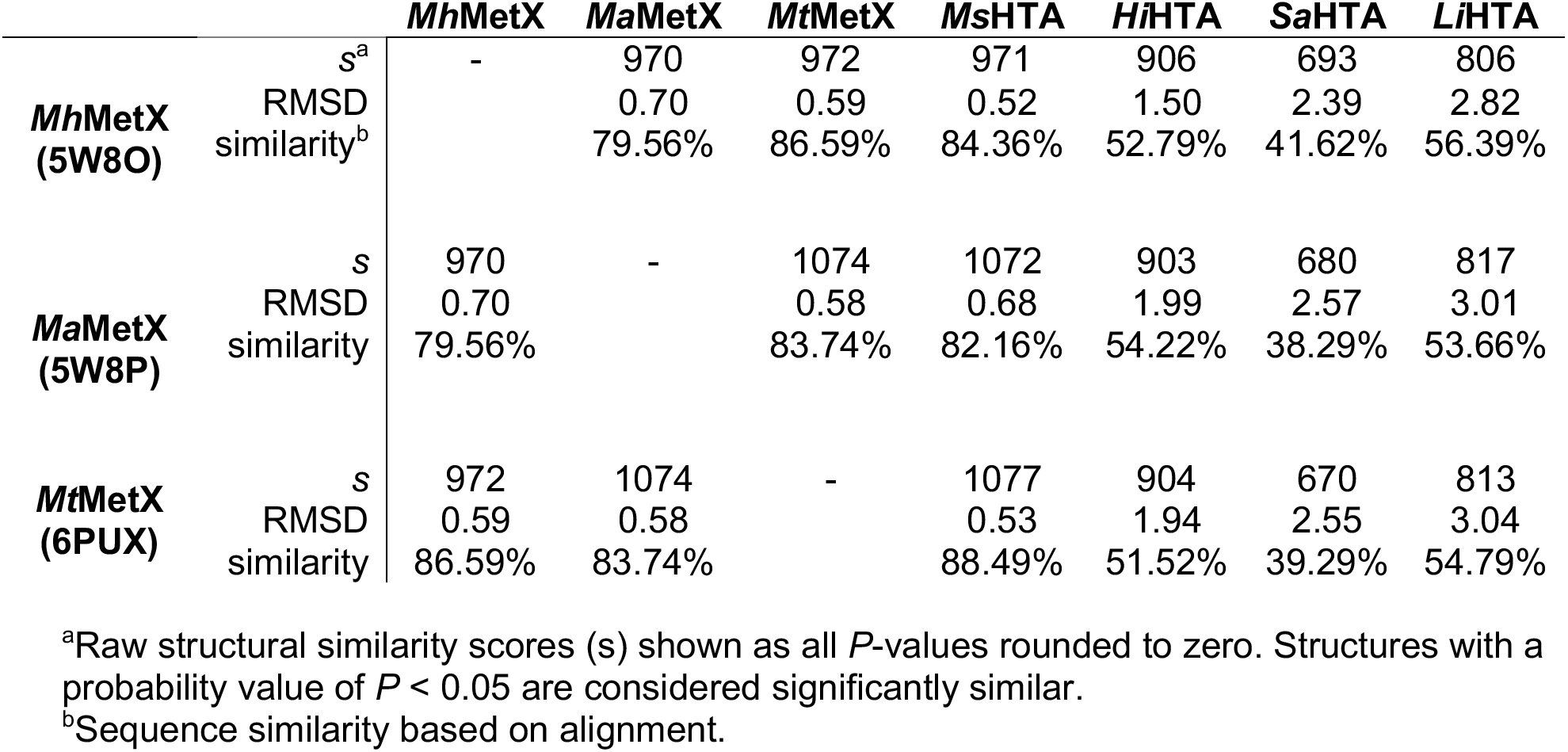
FATCAT pairwise flexible alignment of MetX and HTA structures.

### Differential scanning fluorimetry (DSF)

The assay was carried out using in a range of buffers using 50 mM Tris-HCl pH 6.9-8.5 in 100- 300 mM NaCl with 0.2 mg/mL of *Mt*MetX final concentration. 20 µL of protein was loaded with a 2x final concentration of SYPRO Orange dye (Thermo Fisher) and run on a CFX96 Touch qPCR system (Bio-Rad). A linear thermal ramp of 1°C/min; 20°C - 90°C run with an excitation wavelength of 512–535 nm and a detection wavelength of 560–580 nm. T_m_ was calculated as the minimum of the first derivative plot of the unfolding transition in the CFX Maestro software.

### Structural Comparison and Fragment-based Hot Spot Detection

The Flexible structure alignment by chaining aligned fragment pairs allowing twists (FATCAT) server (http://fatcat.burnham.org) was used to compare the different available HTA structures. All pairwise alignments were done using the flexible alignment model with chain A. The database search for close homologs was performed with a *P*-value of 0.05. The FTMap server (http://ftmap.bu.edu) was used to map chain A of the *Mt*MetX structure and assay ligandability. Cluster strength was determined by the number of probes in each consensus sites. The same run was also performed on the FTFlex server (https://ftflex.bu.edu) to assay whether or not sidechain flexibility would significantly alter the results, but observed trends in cluster locations were not significantly different between both servers.

The *Mt*MetX substrate models were created by using a *Ms*HTA structures, which have been solved in the presence of acetyl-CoA (PDB entries 6IOH and 6IOI)^24^. Monomers from each structure positioned using rigid-body alignment using Chimera’s MatchMaker^49^. Energy minimization on the hybrid structure was then performed using the Molecular Modeling Toolkit and Dock Prep with the AMBER ff14SB forcefield.^50^

## Data Availability

Atomic coordinates and structure factors of the reported crystal structures have been deposited to the Protein Data Bank with accession codes 5W8O (*Mh*MetX), 5W8P (*Ma*MetX), and 6PUX (*Mt*MetX).

## Acknowledgments

The work in this study was supported by an Institutional Development Award (IDeA) from the National Institute of General Medical Sciences of the National Institutes of Health grants P20GM103486 and P30GM110787 to K.V.K. E.S.R. was supported by the National Science Foundation Research Experiences for Undergraduates (REU) grant 1358627. E.S.R. is the American Chemical Society Scholar. C.W.K. was supported by a Howard Hughes Medical Institute undergraduate science education grant to Georgetown College and the Georgetown College Science Honors Program. Use of the Stanford Synchrotron Radiation Lightsource, SLAC National Accelerator Laboratory, is supported by the U.S. Department of Energy, Office of Science, Office of Basic Energy Sciences under Contract No. DE-AC02-76SF00515. The SSRL Structural Molecular Biology Program is supported by the DOE Office of Biological and Environmental Research and by the National Institutes of Health, National Institute of General Medical Sciences (P41GM103393). Use of SER-CAT is supported by its member institutions (see www.ser-cat.org/members.html), and equipment grants (S10_RR25528 and S10_RR028976) from the National Institutes of Health. Use of the Advanced Photon Source was supported by the U. S. Department of Energy, Office of Science, Office of Basic Energy Sciences, contract W-31-109-Eng-38. The contents of this publication are solely the responsibility of the authors and do not necessarily represent the official views of NIGMS or NIH.

## Author Contributions

C.T.C. and K.V.K. planned the experiments. C.T.C., E.S.R., R.W.R., J.L., C.W.K., and K.V.K. performed the experiments. C.T.C. and K.V.K. analyzed the data and wrote the manuscript. All other authors reviewed the manuscript.

